# Estimates of Fall-run Chinook Salmon Escapement in Two San Joaquin River Tributaries from Device-based and Survey-based Methods

**DOI:** 10.1101/2023.10.17.562747

**Authors:** Tyler J. Pilger, Emily Jonagan, Matthew L. Peterson, Jason Guignard, Doug Demko, Andrea Fuller

**Author notes:** Corresponding Author: T.J. Pilger, FISHBIO, 180 E. Fourth Street, Suite 160, Chico, CA 95928, USA.

## Abstract

Monitoring the escapement (i.e., spawner abundance) of adult salmonids is a fundamental component of fisheries stock assessment and evaluating recovery goals. Whereas multiple methods exist for estimating escapement, they exhibit trade-offs between degree of accuracy and implementation expenses and effort. Various methods are used throughout the Central Valley to estimate escapement of the four runs of Chinook Salmon (*Oncorhynchus tshawytscha*), but few rivers have more than one escapement monitoring program. Fall-run Chinook Salmon escapement to the Stanislaus and Tuolumne rivers is estimated by two independent monitoring programs on each river. Since 1952, California Department of Fish and Wildlife has performed carcass surveys that provide an index of escapement. Beginning in 2003 (Stanislaus River) and 2009 (Tuolumne River), private contractors have seasonally installed and operated weirs with fish counting devices to enumerate returning adults. We used the overlapping time series to evaluate the relationship between estimates from carcass surveys and the counting weirs. With few exceptions, estimates from counting adult salmon at the weirs as they entered the spawning grounds were greater than estimates from carcass surveys on both rivers. Escapement estimates from the Stanislaus weir were on average 1.7 (SE = 0.07) times greater than estimates from carcass surveys, while estimates at the Tuolumne weir were 2.5 (SE = 0.17) times greater than estimates from carcass surveys. Estimates from both methods were highly correlated, with *r*^2^ values > 96% for both rivers. The high degree of covariation between methods indicates escapement estimates are robust and can be used in population models that incorporate additional data sources, such as estimates of juvenile production. Having multiple sources of data on fall-run Chinook Salmon will be increasingly useful for guiding and evaluating management actions in these heavily managed rivers.

## INTRODUCTION

Pacific salmonids are some of the most intensively managed fish species in the world due to their economic and cultural value, and because the rivers and streams they inhabit are heavily developed for irrigation, hydroelectric generation, domestic water use, and flood control (Nehlsen et al. 1991; Fisher 1994). For salmonids, estimating spawner escapement (i.e., adults that escape the fishery and return to spawn) is foundational to many management objectives. Escapement monitoring is used to evaluate restoration practices, quantify hatchery contribution to naturally produced stocks, manage for sustainable harvest, and ultimately assess the recovery status of listed threatened and endangered species (Knudsen 1999; Bergman et al. 2012). It also provides a critical data source for salmonid life-cycle models, which are an increasingly used tool to evaluate population-level effects of management actions (e.g., Hendrix et al. 2017; Cordoleani et al. 2020).

Numerous field methods are available to collect abundance data on adult salmonids, such as spawner surveys from aircraft, boat, or on foot; redd and carcass surveys; mark-recapture; and passage counts (Cousens et al. 1982; Johnson et al. 2007). Different field methods produce different data types, and much effort has been spent identifying appropriate statistical frameworks to estimate escapement from various data sources (e.g., Johnson et al. 2007; Parsons and Skalski 2010). Methodologies exhibit different strengths and weaknesses depending on management objectives and logistical or institutional constraints (Cousens et al. 1982; Johnson et al. 2007; Hyun et al. 2012). For example, data from aerial surveys are only appropriate as an index of spawning (Jones et al. 2007), but this may be sufficient for extremely abundant and widespread species, such as pink salmon (*Oncorhynchus gorbuscha*; Bue et al. 1998). Data from carcass and redd surveys are similarly best suited as an index of escapement due to inherent biases related to data collection (e.g., carcasses lost to scavengers and decay [Quinn et al. 2009]) or assumptions about the data (e.g., number of redds per female [Murdoch et al. 2009] and male-to-female ratios [Murdoch et al. 2010]). However, these surveys are more feasible when field personnel or funding is limited. In contrast, counting fish as they pass through a reference point, weir, or ladder, is potentially the most accurate and precise method for estimating absolute escapement, although certain circumstances must be met (Parsons and Skalski 2010). Infrastructure or expensive equipment is required to conduct passage counts, and river discharge and debris loads must remain low for weirs to remain operational during a migration season (Jasper et al. 2018). In addition, estimation accuracy decreases when fish passages per hour are exceedingly high (>1,500 fish per hour; Shardlow and Hyatt 2004), or if morphologically similar species are present and are mistaken for target species (Bergman et al. 2012).

Monitoring Chinook Salmon (*O. tshawytscha*) escapement in California’s Central Valley began in the 1940s and 1950s and has incorporated multiple methods, including redd counts, carcass surveys, and passage counts (Pipal 2005; Low 2007; Bergman et al. 2012). The earliest approaches relied on visual survey methods and later transitioned to mark-recapture of carcasses. Although direct counts have been implemented by Central Valley hatcheries, the first passage counts using an Alaskan-style resistance board weir with a counting device began on the Stanislaus River in 2003. Passage counts are increasingly being used to enumerate adult Chinook Salmon and Steelhead (*O. mykiss*) in the Central Valley (Bergman et al. 2012, Killam 2022). Fall-run Chinook Salmon escapement estimates have been produced from both carcass surveys and passage counts on the Stanislaus River since 2003, and on the Tuolumne River since 2009. Few other rivers in the Central Valley have had multiple escapement methods performed over as many years as these tributaries to the San Joaquin River. Whereas estimates from the two approaches are not directly comparable because one is based on a likelihood mark-recapture estimator and the other is based on observational counts, evaluating the relationship between them can be informative for understanding the robustness of both types of data. Our goal was simply to quantify the covariation between annual estimates from both methods.

## MATERIALS AND METHODS

### Weir and Fish Counting Device

On the Stanislaus and Tuolumne rivers, Alaskan-style resistance board weirs fitted with an infrared and visual fish counting device (Vaki Riverwatcher, Merck & Co., Rahway, NJ, USA) were deployed seasonally to provide passage counts of migrating adult fall-run Chinook Salmon and Steelhead. We used data on the Stanislaus River from 2003 to 2021 and on the Tuolumne River from 2009 to 2021. In most years, the weirs were installed in September, prior to adult fall-run Chinook Salmon migration, and operated through the end of the spawning season, into January or later for monitoring Steelhead. The Stanislaus weir was located 52.6 km upstream of the mouth of the Stanislaus River, and the Tuolumne weir was located 39.4 km upstream of its confluence the San Joaquin River (Figure 1). Both weir locations were selected for ease of river access and for optimal substrate criteria for anchoring the weirs (i.e., uniform channel cross-section with gradual sloping banks and intermixed sand and gravel substrate). Locations were also at the downstream extent of known salmon spawning sites; however, some spawning has been infrequently observed below the weirs (Peterson et al. 2020, J. Guignard, pers. observation). These weirs can only be operated at flows under 1,500–2,000 cubic feet per second (cfs; Stanislaus) or 1,200–1,500 cfs (Tuolumne), depending on the amount of debris. Instantaneous turbidity (Nephelometric Turbidity Units [NTU]) was measured from water samples collected during routine daily checks of both weirs using a turbidity meter (Model 2020e, LaMotte Company, Chestertown, Maryland).

**Figure 1.**
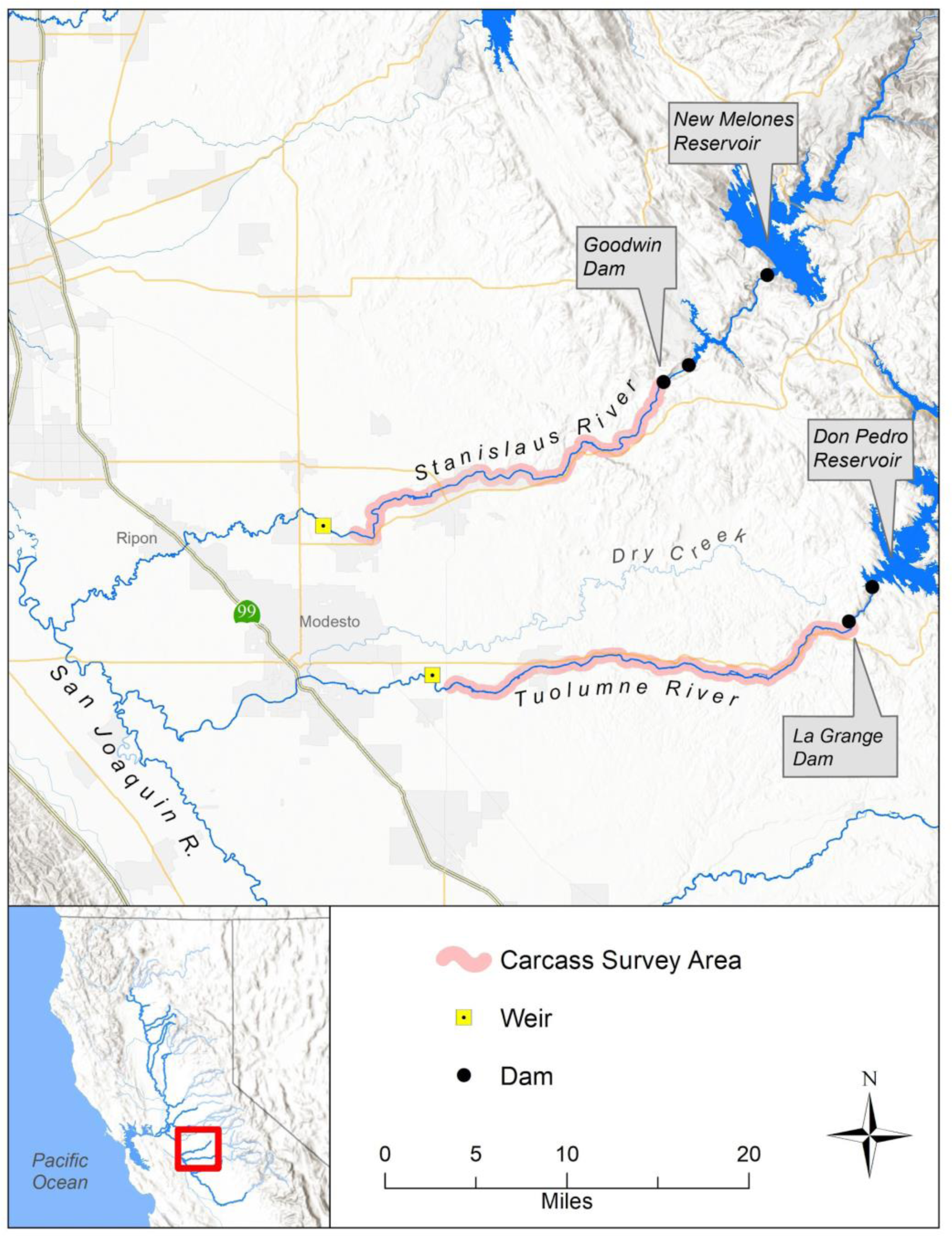
Escapement monitoring on the Stanislaus and Tuolumne rivers. Weirs with fish counting devices are located at the downstream extent of known spawning areas. Carcass surveys are performed throughout the known spawning areas.

The counting device is installed in the weir passage chute and is triggered when a fish or other object enters and crosses through a network of infrared beams emitted from diodes. A scanner plate opposite of the diodes produces a silhouette image of the fish (Shardlow and Hyatt 2004). The device also includes a camera, which takes a series of still photographs or a video clip of the passing fish as the infrared scanner is triggered. Silhouettes and images are only taken and stored if the object detected crossing the scanner beams has a depth that exceeds 40 mm. Additional data recorded include date, time, passage direction (+1 for upstream and -1 for downstream), body depth, and speed passing through the scanner. Occasionally, an additional underwater video system was used as a back-up during prolonged periods (> 2 days) if the counting device malfunctioned. In the office, two experienced reviewers independently examine silhouettes and photographs of each passage using Winari Software (version 4.43, Merck & Co., Rahway, NJ, USA). Silhouettes and photographs were evaluated for image quality and used to identify the species of all individuals counted. Identifications and counts were compared between reviewers, who come to a consensus for any passage record discrepancies.

Passage counts were aggregated into 1-hour time steps for escapement estimates. We used a customized R script (R Core Team 2022) to aggregate counts and, if necessary, impute counts during missing time periods when the device counter was non-operational with no back-up, or when the weir was compromised due to submerged panels (Appendix A of Bergman et al. 2012). Daily counts were imputed via a generalized additive model (GAM) fit to the observed daily counts. Daily passage counts were the sum of observed and any imputed counts. Annual escapement was estimated as the sum of daily counts and we used a bootstrap routine (1,000 iterations) to produce 95% confidence intervals for annual escapement. For the bootstrap, hourly counts within each day were randomly selected with replacement and a GAM was refitted to impute counts for days with missing time periods. The sums of imputed counts for each iteration were used as a distribution of missed counts over the entire season. On days when the device was operating for the full 24-hour period (i.e., no missing time periods), we assumed that detection was perfect and thus no error was associated with counts on those days.

### Carcass Survey Escapement Estimates

Biologists from California Department of Fish and Wildlife (CDFW) have performed mark-recapture carcass surveys to estimate escapement on both rivers since 1971. Surveys were conducted weekly, typically from the first week of October into January. Drift boats were used to access much of each river, where carcasses were collected, scored for their level of decay, and tagged on the lower jaw with a metal disc tag for recapture in subsequent weeks. In reaches that cannot be accessed by drift boat, biologists performed walking surveys to enumerate carcasses. Carcasses not tagged were chopped to prevent double counting. Through the years, several mark-recapture abundance estimators have been used (Bergman et al. 2012), including Jolly-Seber (Jolly 1965; Seber 1965) and Cormack-Jolly-Seber (Cormack 1964) models, as well as superpopulation models (Crosbie and Manly 1985; Schwarz et al. 1993). For river sections where mark-recapture could not be performed, expansion methods were used and the results added to mark-recapture estimates. Details on specific methods for each river can be found in Bergman et al. (2012). We obtained annual fall-run Chinook Salmon escapement estimates for the Stanislaus (2003 – 2021) and Tuolumne (2009 – 2021) rivers from GrandTab (CDFW 2022), an annual compilation of salmon population estimates by CDFW Fisheries Branch Anadromous Resource Assessment Unit.

### Data Analysis

Both weirs were frequently operated past the fall-run Chinook Salmon spawning season for Steelhead monitoring; therefore, to ensure a consistent time frame with the carcass survey estimates, we only included salmon passages from September 1 to December 31 each year. This time frame encompasses more than 95% of cumulative seasonal passage (Peterson et al. 2017). We compared estimates from both methods using simple linear models in R. Estimates from years when the weirs were operated for less than 75% (92 days) of the season, or that had greater than 5% imputed hourly time periods, were excluded from analyses.

## RESULTS

### River Conditions and Weir Operation

On the Stanislaus River, lowest mean seasonal discharge occurred in 2008 (265 cfs, range: 106–801 cfs) and highest mean seasonal discharge occurred in 2011 (1,182 cfs, 367–2,510 cfs). In 2011, discharge exceeded weir operational limits from September through October, delaying installation until November 8 that year. In December of 2005 and 2019, discharge increased to 4,830 cfs and 4,040 cfs, respectively, due to flood control release flows from New Melones Reservoir, necessitating removal of the weir prior to December 31. Turbidity on the Stanislaus was generally low, with 95% of daily instantaneous measurements being less than 10 NTU. However, turbidity greater than 100 NTU was observed in 2012, 2018, and 2021 (Table 1). In 19 years, there were 73 instances when the device malfunctioned for a few hours, and only 5 instances of more than 24 hours elapsing without a backup camera in place. Occasionally, one or more weir panels were submerged upon daily checks, which could allow fish to bypass the counter, and this ranged from 0 to 30 instances per year. The annual percentage of hours for which counts were imputed ranged from 0.0% (2005) to 7.9% (2003). In 2003, there were two multi-day periods during which three panels were removed from the weir (Nov 25 to Dec 1 and Dec 3 to Dec 5), allowing fish to bypass the counter.

**Table 1.** Summary of weir operation data and escapement estimates from the Stanislaus River weir and GrandTab from 2003 to 2021.

| Year | Start Date–<br>End Date | Net<br>Upstream<br>Count<sup>a</sup> | Estimate<br>(95% CI) | GrandTab<sup>b</sup> | Turbidity<br>(NTU)<br>Median (Range) | Percent<br>Missing<sup>c</sup> | Percent<br>Adipose<br>Clipped<sup>d</sup> |
| --- | --- | --- | --- | --- | --- | --- | --- |
| 2003 | 9/3 – 12/31 | 4,319 | 4,901 (4,495 – 5,963) | 5,902 | 1.3 (0.6 – 10.8) | 7.9 | NA |
| 2004 | 9/10 – 12/31 | 4,448 | 4,452 (4,452 – 4,480) | 4,015 | 1.1 (0.6 – 33.0) | 1.1 | 4.5 |
| 2005 | 9/8 – 12/19 | 4,121 | 4,121<sup>e</sup> | 1,427 | 1.3 (0.1 – 4.7) | 0.0 | 2.0 |
| 2006 | 9/8 – 12/31 | 3,019 | 3,023 (3,016 – 3,081) | 1,923 | 0.7 (0.1 – 4.7) | 0.4 | 1.7 |
| 2007 | 9/22 – 12/31 | 438 | 478 (466 – 553) | 443 | 0.8 (0.3 – 4.4) | 2.1 | 3.0 |
| 2008 | 9/9 – 12/31 | 915 | 966 (966 – 1,050) | 865 | 0.6 (0.2 – 31.0) | 0.0 | 4.8 |
| 2009 | 9/9 – 12/31 | 1,256 | 1,371 (1,291 – 1,553) | 595 | 0.9 (0.1 – 5.7) | 1.6 | 11.4 |
| 2010 | 9/7 – 12/31 | 1,346 | 1,476 (1,415 – 1,692) | 1,086 | 1.6 (0.1 – 43.2) | 2.6 | 26.8 |
| 2011 | 11/8 – 12/31 | 757 | 908 (816 – 1,132) | 1,309 | 0.7 (0.1 – 1.6) | 2.5 | 87.8 |
| 2012 | 9/11 – 12/31 | 7,065 | 7,852 (7,753 – 9,912) | 4,006 | 1.9 (0.6 – 106.0) | 2.3 | 67.9 |
| 2013 | 9/3 – 12/31 | 5,428 | 5,537 (5,488 – 5,695) | 2,845 | 1.4 (0.4 – 4.1) | 0.8 | 23.5 |
| 2014 | 9/5 – 12/31 | 5,508 | 5,561 (5,554 – 5,643) | 3,060 | 2.0 (0.5 – 62.3) | 0.7 | 22.4 |
| 2015 | 9/15 – 12/31 | 12,617 | 12,662 (12,655 – 12,726) | 6,136 | 1.7 (0.4 – 4.1) | 0.2 | 26.2 |
| 2016 | 9/8 – 12/31 | 14,369 | 14,966 (14,677 – 15,809) | 9,330 | 1.7 (0.8 – 15.3) | 0.2 | 27.1 |
| 2017 | 9/22 – 12/31 | 8,500 | 9,524 (9,411 – 10,854) | 5,655 | 2.1 (0.6 – 5.1) | 3.6 | 30.5 |
| 2018 | 9/5 – 12/31 | 4,774 | 5,080 (4,966 – 5,619) | 2,387 | 1.6 (0.6 – 126.0) | 1.3 | 26.2 |
| 2019 | 8/29 – 12/3 | 2,594 | 2,821 (2,766 – 3,149) | 1,482 | 1.4 (0.4 – 5.1) | 1.7 | 21.3 |
| 2020 | 9/10 – 12/31 | 1,903 | 2,145 (2,062 – 2,487) | 558 | 1.6 (0.2 – 6.0) | 1.6 | 20.0 |
| 2021 | 9/8 – 12/31 | 6,020 | 6,808 (6,281 – 7,927) | 4,344 | 2.0 (0.6 – 105.0) | 1.9 | 23.7 |

a Net upstream count does not include imputed data from missing time periods
b GrandTab estimates from 2009 to 2021 are preliminary
c Percent of 1-hour time periods requiring GAM imputed counts
d Status of adipose was determined from weir passages, status was not determined in 2003
e No counts were imputed, therefore no confidence interval was calculated

On the Tuolumne River, the lowest mean seasonal discharge occurred in 2015 (137 cfs, 78–169 cfs) and the highest mean seasonal discharge occurred in 2010 (1,354 cfs, 304–5,350 cfs). In 2010, discharge exceeded operational limits in December due to flood control releases from Don Pedro Reservoir, necessitating removal of the weir prior to December 31. Similar to the Stanislaus, the Tuolumne River experiences low turbidity levels, with 99% of daily instantaneous measurements being less than 10 NTU. Unlike the Stanislaus, no extreme turbidity events were observed, and all measurements were less than 30 NTU (Table 2). In 13 years, there were 28 instances when the device malfunctioned for a few hours, and only 2 instances of more than 24 hours elapsing without a backup camera in place. Instances of submerged weir panels during daily checks ranged from 0 to 19 per year. The annual percentage of hours for which counts were imputed ranged from 0.1% (2018) to 4.5% (2010).

**Table 2.**
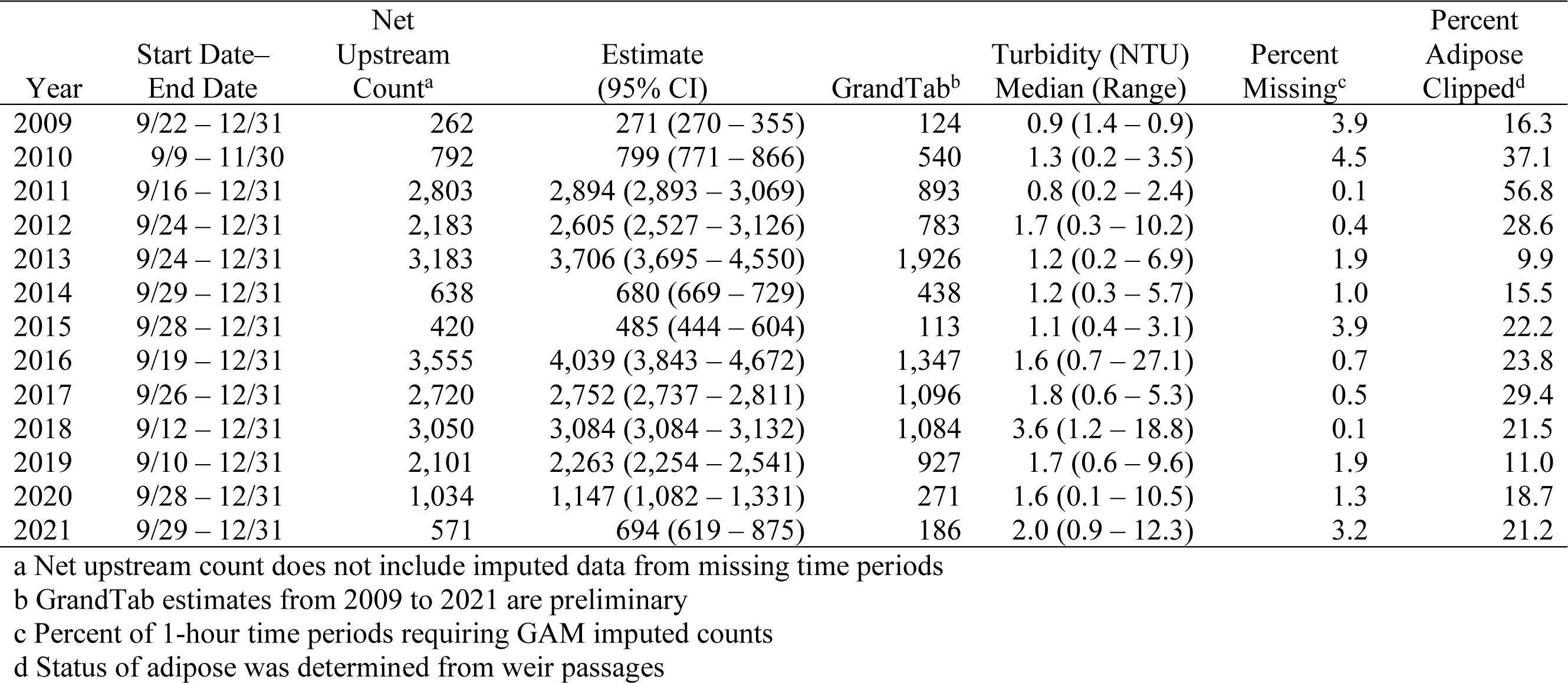
Summary of weir operation data, escapement estimates from the Tuolumne weir and GrandTab from 2009 to 2021.

### Comparison of Escapement Estimates

On the Stanislaus River, carcass survey estimates were greater than weir estimates in only two years, 2003 and 2011, when weir operation was incomplete (Figure 2A). Both methods documented the lowest escapement in 2007 (weir = 478 spawners, carcass = 443 spawners) and greatest escapement in 2016 (weir = 14,966 spawners, carcass = 9,330 spawners). Tuolumne River weir escapement estimates were greater than carcass survey estimates in all years, despite the weir not operating for the complete season in 2010 (Figure 3A). Unlike for the Stanislaus River, the two methods estimated the lowest and highest escapement in differing years for the Tuolumne River. The lowest escapement documented at the Tuolumne weir occurred in 2007 (271 spawners), while the lowest estimate from carcass surveys occurred in 2015 (113 spawners). The highest escapement at the weir was in 2016 (4,039 spawners), but the highest escapement from carcass surveys was in 2013 (1,926 spawners).

**Figure 2.**
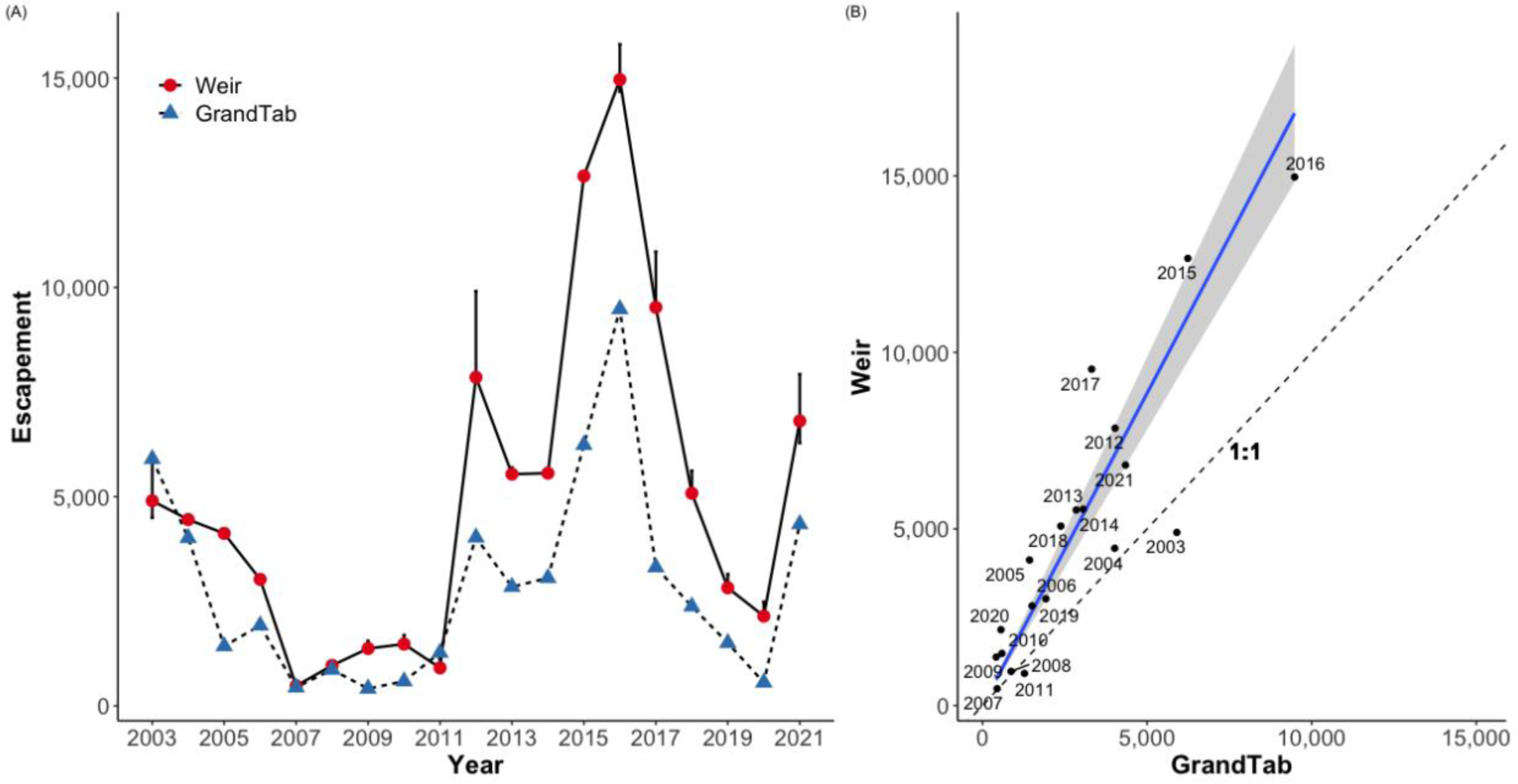
Annual estimates of fall-run Chinook Salmon spawning escapement to the Stanislaus River from 2003 to 2021 (A). Error bars presented on estimates from weir counts (circles) represent bootstrapped 95% confidence intervals. The linear regression (B) excludes estimates from 2003 due to high percentage of imputed hourly counts and partial monitoring year 2011.

**Figure 3.**
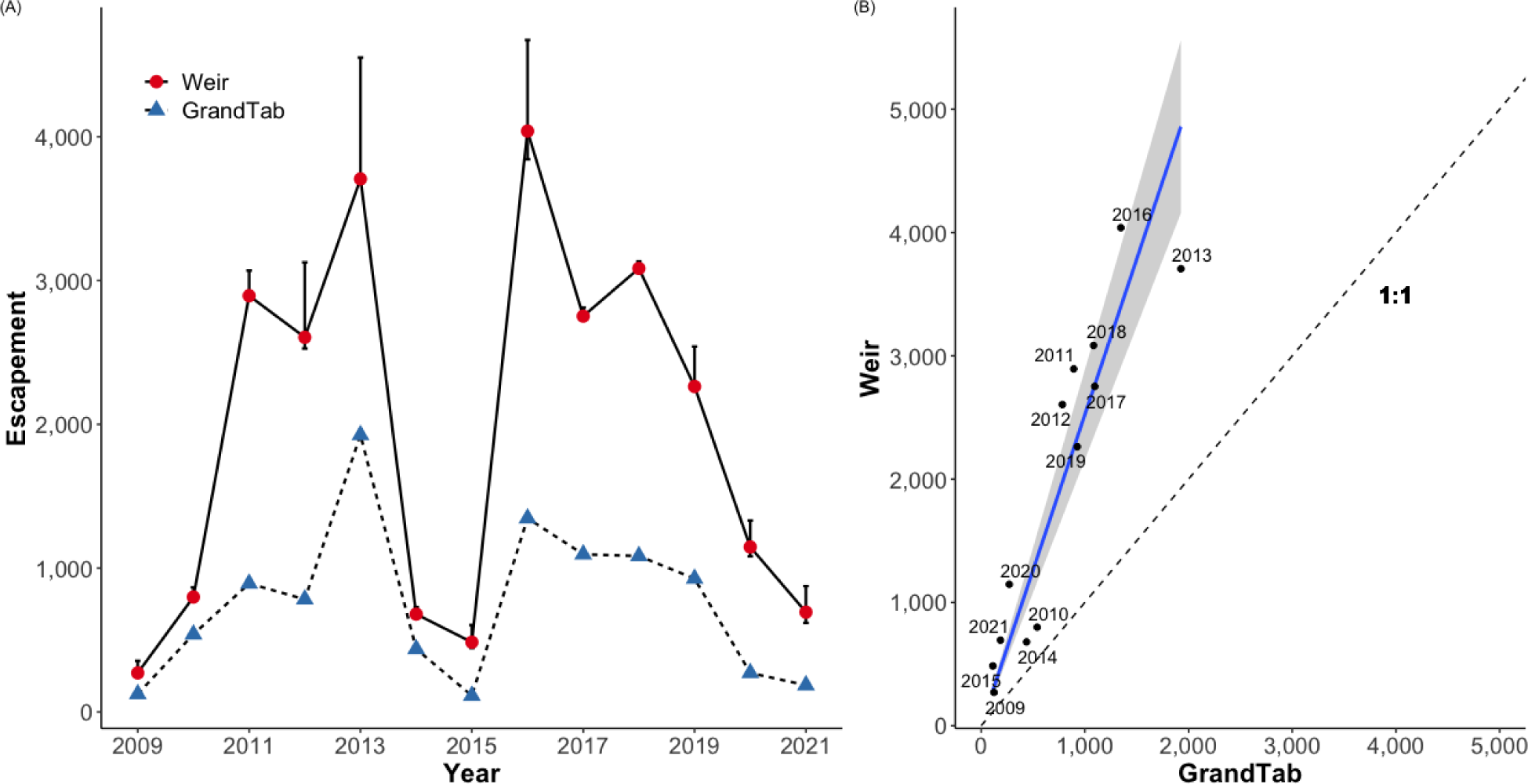
Annual estimates of fall-run Chinook Salmon spawning escapement to the Tuolumne River from 2009 to 2021 (A). Error bars presented on estimates from weir counts (circles) represent bootstrapped 95% confidence intervals. The linear regression (B) excludes estimates from partial monitoring year 2010.

In both rivers, estimates from both methods were highly correlated, but deviated from a 1:1 relationship (Table 3, Figures 2B, 3B). Initial linear models to describe the relationships between methods indicated that the intercepts were not significantly different from zero.

**Table 3.** Coefficients for slope-only linear models comparing weir escapement against carcass survey escapement.

| River | Slope | SE | $r^2$ | p-value | $df$ |
| --- | --- | --- | --- | --- | --- |
| Stanislaus | 1.72 | 0.07 | 0.97 | < 0.001 | 16 |
| Tuolumne | 2.52 | 0.17 | 0.95 | < 0.001 | 11 |

Therefore, we report slope-only models by forcing the intercepts through zero (Table 2). On the Stanislaus, escapement at the weir was on average 1.7 (SE = 0.07) times higher than carcass survey estimates, excluding years 2003 (> 5% imputed hourly counts) and 2011 (incomplete season). The Tuolumne weir escapement was on average 2.5 (SE = 0.17) times higher than carcass survey estimates, excluding the partial year 2010.

## DISCUSSION

For Central Valley Chinook Salmon populations, it is uncommon to have escapement estimates from two independent monitoring programs operating in tandem for as many years as the Stanislaus and Tuolumne rivers (19 and 13 years, respectively). These data sets allowed us to examine annual differences and covariation between estimates of fall-run Chinook Salmon escapement derived from carcass surveys and fish counting devices. The high degree of covariation between these estimates during complete monitoring years means that the index of escapement could be converted to an estimate of absolute abundance using the statistical relationship between them. However, we advise against applying these relationships to historical escapement estimates, especially those generated prior to implementation of mark-recapture methods, because historical abundances were often much greater and would require extrapolation beyond the range of escapement observed since weir monitoring began. Nevertheless, this is an important consideration because the time series of estimates from the carcass surveys are much longer than the weir time series. Additionally, both rivers have long-term juvenile monitoring programs that have operated longer than weir monitoring (e.g., Pilger et al. 2019). Robust data on adult escapement and juvenile production can be used in population models to better understand how changes in environmental conditions and flow management will impact these populations (e.g., Friedman et al. 2019).

Although passage counts at the weirs provide a measure of absolute escapement, these data are not without error (Bergman et al. 2012). Counting devices can underestimate true escapement if individuals are missed due to device malfunction or if extraneous factors, such as turbidity, prevent the device from triggering. Device malfunctions did occur on both rivers, but typically lasted over the span of hours and only rarely days. During time periods when the device was malfunctioning, or when we suspected the weir was compromised (i.e., weir panels were submerged), we used a GAM to impute potentially missed individuals as recommended in Bergman et al. (2012). The counting devices are also known to function with high accuracy over the range of turbidity levels observed in these rivers (Baumgartner et al. 2012). Escapement can also be underestimated if individuals by-pass the device swim channel through holes that form under or to the sides of the weir, or by going over the weir when panels are forced under the water surface (usually by high debris loads). The weirs were inspected daily to ensure no holes formed along the substrate rail or around the sides, and to clean accumulated debris off the panels. During high flows when debris loads can increase, weirs were continuously monitored to keep debris from submerging the panels. Weir panels did get submerged, but on an infrequent basis. Counters can also experience error at high passage rates. Shardlow and Hyatt (2004) showed that accuracy of a similar infrared device is greater than 95% at passage rates lower than 500 fish/hour. Across all years, the maximum passage rate observed was 267 fish/hour at the Stanislaus weir, and 141 fish/hour at the Tuolumne weir, thus passage rates on both rivers were not expected to compromise accuracy.

Some types of error can lead to an over-counting of escapement, including objects other than fish triggering the device, or individuals of a morphologically similar species being counted (e.g., Steelhead). Over-counting can also occur if an individual is counted once passing upstream, then doubles back downstream without getting counted and is counted a second time passing back upstream. Pairing the counting device with photographs of passage events allows us to validate 100% of passages, thus minimizing any potential for estimation bias due to non-fish objects or other fish species triggering the devices (Putt et al. 2022). Steelhead do occur in the Stanislaus and Tuolumne rivers, although infrequently (Eschenroeder et al. 2022), and photographs from the device are sufficient to distinguish them from Chinook Salmon. Lastly, the narrow width of the device swim channel (30 cm) and the sensitivity of the infrared beams make it possible to detect and count individuals passing downstream through the device, which are then subtracted from upstream counts for total passages.

With few exceptions, estimates of absolute abundance from the weirs were greater than the index of escapement estimated from carcass surveys in both rivers. This result is not all that surprising given the inherent bias associated with carcass loss (Quinn et al. 2009; Murdoch et al. 2010). Nevertheless, years such as 2011 on the Stanislaus River demonstrate the importance of having both programs. Flood control releases during the migration season delayed weir monitoring until the majority of the run had passed (Peterson et al. 2017), with a resulting estimate of 908 (816–1,132) spawners. Although high flows also affected early weeks of the carcass surveys that year, few redds and carcasses are usually observed during those weeks at the start of the season (Giudice 2012, unreferenced, see “Notes”). The carcass survey estimate in 2011 was 1,309 spawners. Based on the relationship between the estimation methods, and making some simplifying assumptions about carcass detections, the expected weir escapement would have been 2,251 (2,054–2,448). Due to the number of consecutive missing days in 2011, it was not appropriate to use the GAM approach to estimate counts during that time. A promising alternative to acquire more complete estimates in partial operation years would be to use a hierarchical Bayesian model (Jasper et al. 2018). Such models are capable of sharing information about run-timing and counts using hyperparameters to estimate run curves in years with incomplete monitoring.

The value gained from having two complementary escapement monitoring programs on the Stanislaus and Tuolumne rivers cannot be overstated. In addition to a multi-decadal time series index of fall-run Chinook Salmon escapement, carcass surveys allow for collection of biological samples, such as tissue for genetic assays (Garza et al. 2008), coded wire tags to determine hatchery origin and straying (Huber and Carlson 2015; Sturrock et al. 2019), scale samples for age analysis, and otoliths to track individual life histories (Sturrock et al. 2015, Sturrock et al. 2020). Operation of a weir and fish counting device is not only a non-invasive method for acquiring accurate estimates of the number of adults returning to spawn, but also provides fine-scale data on seasonal migrations. Hourly data can identify the time of day when adults are more actively migrating upstream and daily data can be used to assess how management actions affect seasonal migration patterns (Peterson et al. 2017). Both programs collect data to estimate spawner sex ratios, body length, and proportion whose adipose fin was clipped (i.e., a mark indicating hatchery origin). Using data from both programs may allow for refined understanding of spawner composition. For example, Central Valley hatcheries use constant fractional marking for fall-run Chinook Salmon, and estimating hatchery contribution to in-river spawners requires both the proportion of adipose fin clipped adults and recoveries of coded wire tags (Mohr and Satterthwaite 2013). Although it is not possible to confirm adipose fin status for 100% of salmon passing through the counting device due to insufficient lighting or body orientation, the proportion of adipose fin clipped individuals observed at the weir is potentially more representative of the spawning population than estimates made from carcasses. This is because the soft adipose tissue decomposes quickly and only a fraction of carcasses are handled due to being lost to scavengers, washed downstream, or never being detected. Thus, data on adipose fin status from the weir could be used to refine estimates of hatchery contribution on these rivers. A first step to this end would be to compare the proportions of adipose fin clipped individuals between monitoring programs.

The Stanislaus and Tuolumne rivers are important sources of natural production of Central Valley fall-run Chinook Salmon, and escapement monitoring is vital to tracking the status of these populations. Complementary escapement monitoring programs are needed because different monitoring methods have tradeoffs associated with the cost and effort of implementation, as well as the precision and accuracy of estimates (Hyun et al. 2012). Flows on these rivers are highly regulated for hydroelectric generation, agricultural and urban water use, and to benefit fish. Management actions such as pulse flows, unimpaired flows, and habitat restoration have been implemented with the goal of benefiting native salmonids; however, the ability to detect changes in naturally variable populations in response to these management actions requires both estimates with high precision and decades of consistent monitoring (Korman and Higgins 1997).

## ACKNOWLEDGEMENTS

The United States Fish and Wildlife Service provided funds for the purchase of the Vaki Riverwatcher and weir components for pilot operation on the Stanislaus River, and long-term operation was funded by Oakdale and South San Joaquin Irrigation Districts. All funding for equipment and long-term operation of the weir and Vaki Riverwatcher on the Tuolumne River was provided by Turlock Irrigation District, Modesto Irrigation District, and the City and County of San Francisco. We thank numerous people who have assisted with these monitoring programs, including FISHBIO staff that constructed, operated, and maintained the weirs over time. Ryan Nielson provided the initial code for device counter analyses. Annual carcass surveys to estimate escapement were performed by California Department of Fish and Wildlife. Use of trade names does not imply endorsement by federal or state agencies or the irrigation districts. Versions of this manuscript were greatly improved by comments from Shaara Ainsley, Erin Loury, two anonymous reviewers, and the editors.

## NOTES

Giudice J. 2012. Stanislaus River Fall Chinook Salmon Escapement Survey 2011 Draft. For United States Bureau of Reclamation. February 2012. 23 pp. Provided by California Department of Fish and Wildlife.

